# Crystal structure of schizorhodopsin reveals mechanism of inward proton pumping

**DOI:** 10.1101/2020.07.28.224907

**Authors:** Akimitsu Higuchi, Wataru Shihoya, Masae Konno, Tatsuya Ikuta, Hideki Kandori, Keiichi Inoue, Osamu Nureki

**Affiliations:** Department of Biological Sciences, Graduate School of Science, The University of Tokyo, Bunkyo, Tokyo 113-0033, Japan; The Institute for Solid State Physics, The University of Tokyo, Kashiwa 277-8581, Japan; PRESTO, Japan Science and Technology Agency, 4-1-8 Honcho, Kawaguchi, Saitama 332-0012, Japan; Department of Life Science and Applied Chemistry, Nagoya Institute of Technology, Showa, Nagoya 466-8555, Japan; OptoBioTechnology Research Center, Nagoya Institute of Technology, Showa, Nagoya 466-8555, Japan

## Abstract

Schizorhodopsins (SzRs), a new rhodopsin family identified in Asgard archaea, are phylogenetically located at an intermediate position between type-1 microbial rhodopsins and heliorhodopsins. SzRs reportedly work as light-driven inward H^+^ pumps, as xenorhodopsin. Here we report the crystal structure of SzR AM_5_00977 at 2.1 Å resolution. The SzR structure superimposes well on that of bacteriorhodopsin rather than heliorhodopsin, suggesting that SzRs are classified with type-1 rhodopsins. The structure-based mutagenesis study demonstrated that the residues N100 and V103 are essential for color tuning in SzRs. The cytoplasmic parts of transmembrane helices 2, 6, and 7 in SzR are shorter than those in the other microbial rhodopsins. Thus, E81 is located near the cytosol, playing a critical role in the inward H^+^ release. We suggested the H^+^ is not metastably trapped in E81 and released through the water-mediated transport network from the retinal Schiff base to the cytosol. Moreover, most residues on the H^+^ transport pathway are not conserved between SzRs and xenorhodopsins, suggesting that they have entirely different inward H^+^ release mechanisms.

## Introduction

Microbial rhodopsins are a large family of heptahelical photoreceptive membrane proteins that use retinal as a chromophore^1^. They are found in diverse microorganisms, such as bacteria, archaea, alga, protists, fungi and giant viruses^2–4^. The retinal chromophore in the microbial rhodopsins undergoes all-*trans* to 13-*cis* isomerization upon light illumination, leading to a photocyclic reaction in which the proteins exert their various biological functions. Ion transporting rhodopsins are the most abundant microbial rhodopsins, and are classified into light-driven ion pumps and light-gated ion channels. Whereas light-driven ion pumps actively transport ions in one direction, light-gated ion channels passively transport them according to the electrochemical potential. Ion transporting rhodopsins are used as important molecular tools in optogenetics, to control neural firing *in vivo*. Microbial rhodopsins evolved independently from animal rhodopsins, which are also retinal-bound heptahelical proteins and a sub-group of G-protein coupled receptors. The third class of rhodopsin, heliorhodopsin (HeR), was recently reported. It has an inverted protein orientation in the membrane, as compared with microbial and animal rhodopsins^5^.

Bacteriorhodopsin (BR) is the first ion pump rhodopsin found in the haloarchaeon^6^, *Halobacterium salinarum*, and it transports protons (H^+^) outward. An inward chloride (Cl^-^) pump, halorhodopsin, was subsequently identified in the same species^7,8^. Although an outward sodium (Na^+^) pump rhodopsin was not found for several decades after the discovery of BR, it was eventually identified in the marine bacterium *Krokinobacter eikastus* in 2013^9^. These ion-pumping rhodopsins hyperpolarize the membrane by their active ion transport against the electrochemical potential of the membrane. However, the bacterial xenorhodopsins (XeRs) reportedly work as light-driven inward H^+^ pumps^10^. Thus, the membrane potentials are not exclusively hyperpolarized via active transport by ion pumping.

Asgard archaea are the closest prokaryotic species to ancestral eukaryotes^11^ and have many genes unique to eukaryotes. Recently, a new microbial rhodopsin group, schizorhodopsin (SzR), was found in the assembled genomes of Asgard archaea and the metagenomic sequences of unknown microbial species^12,13^. A molecular phylogenetic analysis suggested that SzRs are located at an intermediate position between typical microbial rhodopsins, also called “type-1 rhodopsins”^14^, and HeR^5^, and thus they were named “schizo- (meaning “split” in Greek)” rhodopsin. Especially, the transmembrane helix (TM) 3 of SzR is more similar to that of HeR than type-1, whereas TM6 and 7 of SzR and type-1 share many identical residues; e.g., W154, P158, T161, A184, and F191, which are not conserved in HeR^13^. SzRs heterologously expressed in *E. coli* and mammalian cells displayed light-driven inward H^+^ pump activity^13^. As SzRs are phylogenetically distant from XeRs (~18% identity and ~44% similarity), these two rhodopsin families with similar functions have convergently evolved.

In both SzR and XeR, an H^+^ is released from the Schiff base linkage connecting the retinal and a conserved lysine residue (retinal Schiff-base, RSB) in TM7 to the cytoplasmic side. The transiently deprotonated RSB shows a largely blue-shifted absorption peak, and this blue-shifted state was named the M-intermediate. In the case of XeR from the marine bacterium *Parvularcula oceani* (*Po*XeR), the H^+^ is transferred to the cytoplasmic aspartate (*Po*XeR D216, H^+^ acceptor) in TM7, and then released to the cytoplasmic bulk phase^10^. By contrast, the H^+^ acceptor of SzR was considered to be E81 in TM3, since the mutation of E81 to glutamine abolished the inward H^+^ transport^13^. However, the H^+^ is not metastably trapped in E81, probably for a kinetic reason: the rate of H^+^ release from E81 to the cytoplasmic bulk phase might be faster than that of H^+^ transfer from RSB to E81. The reason why SzR and *Po*XeR exhibit different kinetic behaviors in H^+^ release has not been elucidated. Subsequently, another H^+^ is taken up from the extracellular side, and directly transferred from the extracellular bulk phase to the RSB during the M-decay to the initial state.

Recently, a new SzR sub-group, AntR, was identified in metagenomic data obtained from Antarctic freshwater lake samples^15^. Although SzR and AntR share substantial similarity (identity: ~33%; similarity: ~56%) and most of the SzR residues essential for the inward H^+^ pump function are conserved in AntR (e.g., SzR R67, F70, C75, E81, D184, and K188), they have several differences. Notably, while the SzR E81Q mutant cannot transport H^+^, as mentioned above, the H^+^ transport efficiency of AntR E81Q is close to that of AntR wild-type (WT)^15^, suggesting the diversity of H^+^ transport mechanisms. To understand the inward H^+^ pump mechanism of SzR, as well as the similarities and differences between SzR, XeR, and AntR, we present the first 3D-structure of an SzR.

### Functional characterization of SzR4

For the structural analysis, we screened multiple SzRs and identified SzR AM_5_00977 (GenBank accession number: TFG21677.1, hereafter called SzR4) as a promising candidate (Fig. 1a). We purified and crystallized the full-length SzR4, using *in meso* crystallization. Eventually, we determined the 2.1 Å resolution structure of SzR4, by molecular replacement using BR as the search model (Table 1).

**Fig. 1.**
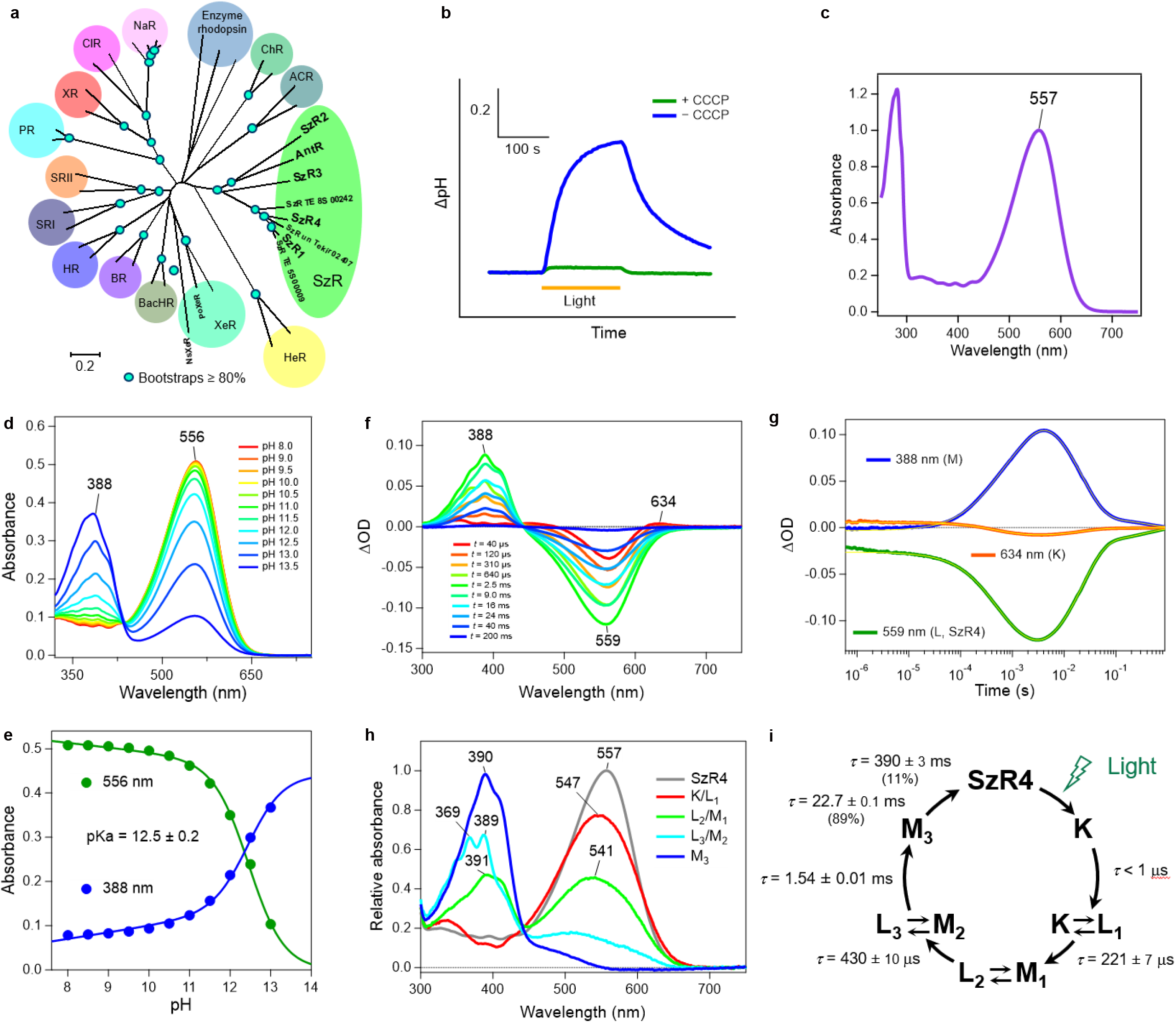
Characterization of molecular properties of SzR4. **a**, Phylogenetic tree of microbial rhodopsins. **b**, H^+^-transport activity assay of SzR4 in *E. coli* cells suspended in 100 mM NaCl. Blue and green lines indicate the results in the absence and presence of CCCP, respectively. **c**, UV-visible absorption spectrum of purified SzR4. **d**, **e**, UV-visible absorption spectra (**d**), and absorption of SzR4 at *λ* = 556 (green circles) and 388 nm (blue circles) at pH 8.0-13.5 (**e**) in 100 mM NaCl 6-mix buffer (citrate, MES, HEPES, MOPS, CHES, CAPS, 10 mM each) containing 0.05% DDM. The solid lines in the latter indicate lines obtained by global fitting with the Henderson– Hasselbalch equation, and the pKa was determined to be 12.5 ± 0.2 (mean ± s.d.). **f**, **g**, Transient absorption spectra of SzR4 (**f**) and time-evolutions at specific wavelengths representing each state (388 nm: the M intermediate; 559 nm: the L intermediate and SzR4; 634 nm: the K intermediate) (**g**) in 100 mM NaCl, 20 mM Tris-HCl, pH 8.0, POPE/POPG (molar ratio 3:1) vesicles with a lipid to protein molar ratio = 50. The thin yellow lines in the latter indicate the lines obtained by global fitting with a multi-exponential function. **h**, **i**, Calculated absolute absorption spectra of the initial state and the photo-intermediates (**h**) and the photocycle (**i**) of SzR4, based on the fitting shown in e and a kinetic model assuming a sequential photocycle. The lifetime (*τ*) of each intermediate is indicated by mean ± s.d. The numbers in parentheses indicate the fraction of the M intermediate decayed with each lifetime in its double-exponential decay.

**Table 1.** Data collection and refinement statistics.

| SzR4 |  |
| --- | --- |
| **Data collection** |  |
| Space group | *C2* |
| Cell dimensions |  |
| *a*, *b*, *c* (Å) | 106, 61.1, 98.8 |
| $\alpha$, $\beta$, $\gamma$ (°) | 90, 99.353, 90 |
| Resolution (Å)* | 48.09 - 2.10 (2.175 - 2.10) |
| *R*<sub>meas</sub>* | 0.3922 (3.369) |
| $\langle I/\sigma(I) \rangle$* | 6.81 (1.18) |
| CC<sub>1/2</sub>* | 0.988 (0.39) |
| Completeness (%)* | 99.56 (98.45) |
| Redundancy* | 19.3 (19.6) |
| **Refinement** |  |
| Resolution (Å) | 49.56-2.10 |
| No. reflections | 36443 |
| *R*<sub>work</sub> / *R*<sub>free</sub> | 0.1976 / 0.2376 |
| No. atoms |  |
| Protein | 4858 |
| Ligand | 780 |
| Water | 144 |
| Averaged *B*-factors (Å<sup>2</sup>) |  |
| Protein | 32.7 |
| Ligand | 39.3 |
| Water | 69.8 |
| R.m.s. deviations from ideal |  |
| Bond lengths (Å) | 0.002 |
| Bond angles (°) | 0.527 |
| Ramachandran plot |  |
| Favored (%) | 98.6 |
| Allowed (%) | 1.4 |
| Outlier (%) | 0 |

\*Values in parentheses are for highest-resolution shell.

We first characterized the biochemical properties of SzR4. The phylogenetic tree of microbial rhodopsins indicated that SzR4 belongs to the SzR family, which is far from xenorhodopsin (XeR), and it is close to the previously characterized SzR1 (Fig. 1a). To investigate the ion transport function of SzR4, we exposed SzR4-expressing *Escherichia coli* to visible light and observed alkalization of the external solvent (Fig. 1b). The alkalization was largely eliminated by the addition of a protonophore, 10 μM CCCP (carbonyl cyanide m-chlorophenylhydrazone), suggesting that SzR4 functions as an inward H^+^ pump as reported previously^13^. The purified SzR4 showed a maximum absorption wavelength (*λ*_max_) at 557 nm, identical to that of SzR1^13^ (Fig. 1c). The absorption peak in the visible wavelength region was decreased at higher pH, and another peak appeared in the UV region (*λ*_max_ = 388 nm) (Fig. 1d, e). The latter represents the deprotonation of the RSB^13^, and its pKa was 12.5 ± 0.2 (mean ± s.d.). This is 1 unit smaller than that of SzR1, suggesting that the protonated RSB is less stabilized in SzR4.

To investigate the photocycle of SzR4, we performed a laser flash photolysis experiment with SzR4 in POPE/POPG vesicles. Transient absorptions representing the accumulations of K, L, and M intermediates were observed, as in the photocycle of SzR1 ^13^(Fig. 1f, g). The sum of five-exponential functions effectively reproduced the time evolution of the transient absorption change of SzR4. The absorption spectra of the initial state and four photo-intermediates (K/L1, L2/M1, L3/M2, and M3) and the photocycle of SzR4 were determined (Fig. 1h, i). The overall photoreaction cycle of SzR4 is similar to that of SzR1^13^. A large accumulation of the M intermediate was observed in the millisecond region, representing the deprotonated state of the RSB. An equilibrium exists between M1 and M2 with L at different equilibrium constants, and it is more biased towards the M for L3/M2 than for L2/M1. Notably, the absolute spectra of the three M states were substantially different. Specifically, the vibrational structure observed in M2 was less pronounced in M1 and M3 (Fig. 1h). This spectral change would originate from a large conformational change of the protein around the retinal chromophore, and be associated with the conversion from the inward opened to outward opened state. The H^+^ release to the cytoplasmic side is finished until the M3 formation, and thus a new H^+^ is taken up from the extracellular side during the M3-decay^13^. A similar spectral change in the vibrational structures between two M states was also reported for XeR from *Nanosalina* (*Ns*XeR)^16^, suggesting that a comparable conformational change also occurs between the H^+^ release and uptake processes in SzR and XeR.

### Overall structure of SzR4

The crystallographic asymmetric unit contains three molecules (mol A, mol B, and mol C) (Extended Data Fig. 1a, b). The overall architectures of these three molecules are almost identical, and thus we focused on the mol A structure. SzR4 consists of 7 TMs and six loops (extracellular loops 1-3 and intracellular loops 1-3) (Fig. 2a, b). Five residues after H199 are disordered in the crystal structure. Extracellular loop 2 (residues G51-Y64) contains a short anti-parallel β strand. All-*trans* retinal is covalently bound to K188, forming the RSB, as in other microbial rhodopsins.

**Fig. 2.**
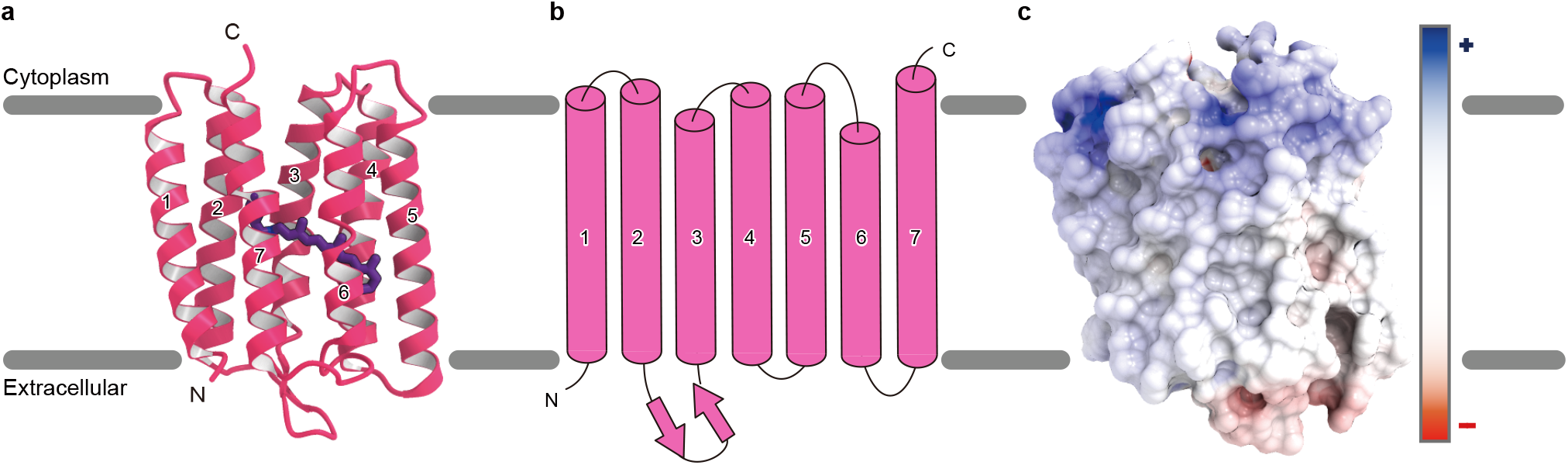
Overall structure of SzR4. **a**, Ribbon diagrams viewed from the membrane plane (right). **b**, Schematic representation of the SzR4 structure. **c**, Electrostatic surface viewed from the membrane plane. Red and blue correspond to potentials of −8 kT e^-1^ and 8 kT e^-1^, respectively.

A previous immunostaining analysis revealed that the C terminus of SzR1 is oriented toward the cytoplasmic side^13^, as in the type-1 rhodopsins, which is opposite to the HeRs. Many positively and negatively charged residues are present on the cytoplasmic and extracellular faces, respectively, in the SzR4 structure (Fig. 2c). This electrostatic surface is consistent with the positive-inside rule^17^, and also supports its topology.

The three SzR4 molecules in the asymmetric unit form a trimer in the crystal structure, in excellent agreement with the previous HS-AFM observation. TM1 and TM2 of one protomer interact with TM4’ and TM5’ of the adjacent protomer, creating the trimer interface (Fig. 3a and Extended Data Fig. 1c). The interface comprises mainly hydrophobic residues (Extended Data Fig. 1d, e), and several hydrogen-bonding interactions are observed on the cytoplasmic side. The residues at the interface are conserved among SzRs (Extended Data Fig. 1f), indicating that SzRs generally function as trimers.

**Fig. 3.**
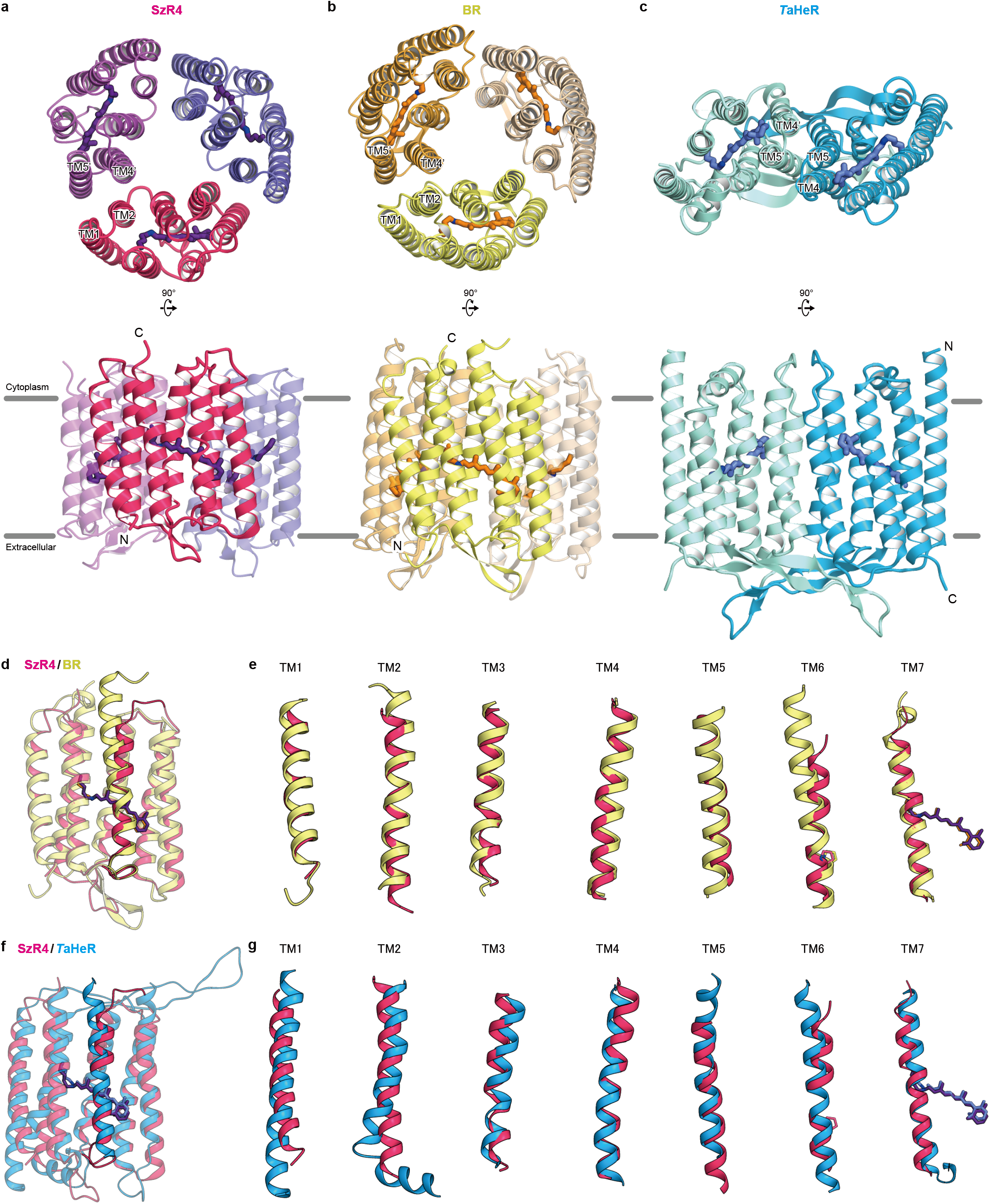
Comparison of SzR4, BR, and *T*aHeR. **a-c**, Monomer and oligomeric structures of SzR4 (**a**), BR (Protein Data Bank (PDB) code: 1M0L) (**b**), and *T*aHeR (PDB code: 6IS6) (**c**), colored magenta, yellow, and dark turquoise. **d**, **e**, Superimpositions of the SzR4 and BR structures. Individual TM helices are shown after superimposition of the two rhodopsins (**e**). **f**, **g,** Superimpositions of the SzR4 and *T*aHeR structures (**f**). Individual TM helices are shown after superimposition of the two rhodopsins (**g**).

To determine whether SzRs are classified as either type-1 rhodopsins or HeRs, we compared the structures of SzR4, BR, and *T*aHeR. SzR4 and BR similarly form trimers, while *T*aHeR forms a dimer (Fig. 3a-c). The SzR4 and BR structures also have the same configuration of TMs forming trimeric binding interfaces, with TM1 and TM2 of one monomer creating a binding interface with TM4 and TM5 of the adjacent monomer. SzR4 and BR also share a common orientation relative to the membrane. Moreover, the monomer structure of SzR4 superimposes well on that of BR (R.M.S.D. = 1.23 Å) (Fig. 3d, e). By contrast, the orientations of SzR4 and *T*aHeR are reversed in the membrane. When the N- and C-termini of SzR4 and *T*aHeR are aligned and their monomeric structures are superimposed, the slope and length of each TM do not overlap well, as compared to BR (R.M.S.D = 2.27 Å) (Fig. 3f, g). Overall, although SzR4 has approximately 20% sequence identity to both BR and HeR, it is structurally more similar to BR. Hence, we suggest that SzRs belong to the type-1 rhodopsins.

We next compared the SzR4 and BR structures in detail. Each TM overlaps relatively well, and their ECL2 similarly contain an anti-parallel β-strand (Fig. 3d). However, there is a striking difference on the cytoplasmic side. The C-terminus of BR contains a short α-helix and is directed toward the center of the protein, while the C-terminus of SzR4 is disordered. Moreover, TM2 and TM6 of SzR4 are shorter than those of BR by one and two α-helical turns, respectively. Notably, the length between the conserved Pro and the cytoplasmic end of TM6 is 13 residues in SzR4, while those in other type-1 rhodopsins are about 21 residues. Thus, the cytoplasmic part of TM6 in SzR4 is the shortest among the microbial rhodopsins (Extended Data Fig. 3a, b). The sequence alignment of SzRs revealed that the shorter length of TM6 is highly conserved (Extended Data Fig. 2), and thus it is a unique structural feature of SzRs.

### Retinal binding site and color tuning mechanism

We next describe the retinal binding pocket in SzR4 (Fig. 4a). D184 forms a direct salt bridge with the RSB and functions as a single counterion, stabilizing the high pKa (12.5, Fig. 1e) of the RSB. The other residues in the retinal binding pocket are mainly hydrophobic. Notably, the aromatic residues Y71 and W154 closely contact the C10-C13 moiety of the retinal from below and above, respectively, allowing the *all-trans* to 13-*cis* isomerization^13^. These residues are completely conserved in 85 SzR homologs (Fig. 4b). The equivalent residues are two tryptophan residues in type-1 rhodopsins, and tyrosine and phenylalanine residues in HeRs. From this viewpoint, SzR is in between BR and HeR. Among the residues constituting the retinal binding pocket, 7 and 3 residues of SzR4 are conserved in BR and *T*aHeR, respectively. Thus, the retinal binding pocket of SzR is similar to that of type-1 rhodopsins, rather than HeRs,

**Fig. 4.**
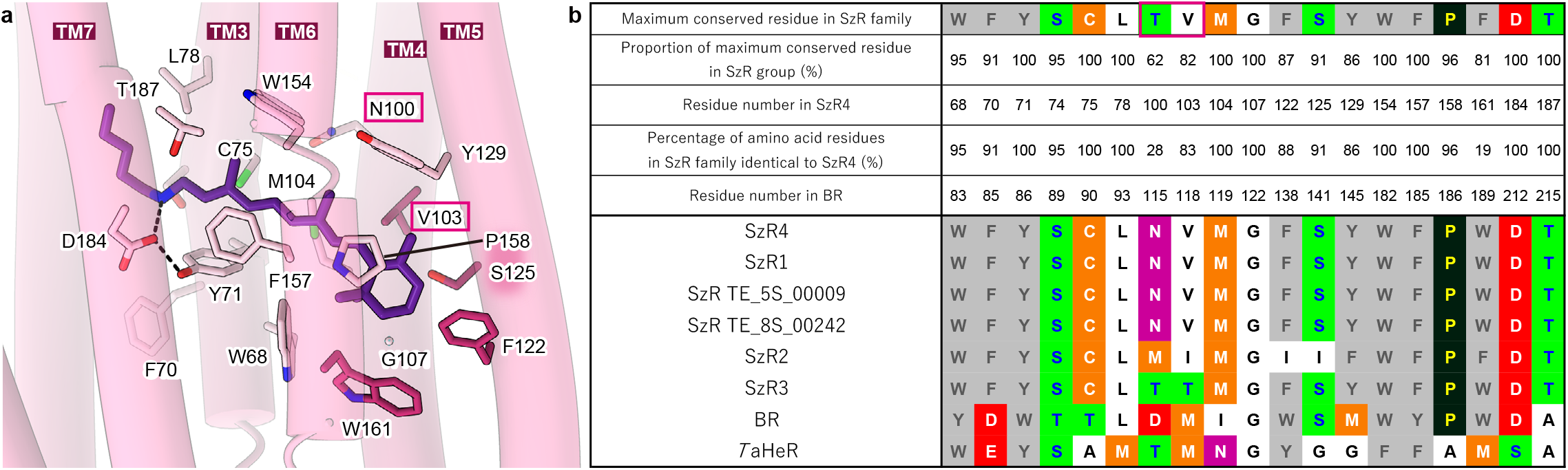
Conservation of retinal binding site. **a**, The retinal chromophore and the residues within 4.5 Å involved in retinal binding. **b**, Maximally conserved residues around retinal and their percentages in SzR family members, with residue numbering according to SzR4. The variations of the amino acid types in six SzRs, BR, and *T*aHeR are shown in the lower part.

The environment around the retinal chromophore is closely associated with the absorption wavelength of rhodopsins. The purified SzR4 displayed the *λ*_max_ at 557 nm, which is identical to that of SzR1, and 555 nm when expressed in *E. coli* cells. Notably, SzR2 and SzR3 showed blue-shifted absorptions at 542 and 540 nm, respectively (Table 2 and Extended Data Fig. 4). To investigate the color tuning mechanism in SzRs, we compared the residues constituting the retinal binding pocket between the six homologs of SzRs (Fig. 4b). These residues are entirely conserved in SzR4, SzR1, SzR TE_5S_00009, and SzR TE_8s_00242. Comparing SzR4, SzR2, and SzR3, the residues around the polyene chain are entirely conserved, whereas those around the β-ionone ring are more diverged. V103, F122, S125, and W161 in SzR4 are replaced in SzR2, and V103 is replaced in SzR3. Moreover, in the vicinity of the β-ionone ring, N100 in SzR4 is replaced with M and T in SzR2 and SzR3, respectively. In BR, D115 is present at the homologous position. BR D115N and D115A showed 2- and 11-nm blue-shifts as compared with the WT, respectively^18,19^, and thus a different amino acid at this position would generate distinct *λ*_max_ values among SzR4, SzR2, and SzR3.

**Table 2.** Absorption maximum positions of SzR4, SzR2, and SzR3 and their mutants. *Λ*_max_: The maximum absorption wavelength Δ*λ*_max_: The difference from the wildtype protein *λ*_max_ values of SzR4 S125Y, V103T, and T187V could not be determined, due to their low expression in *E. coli*.

| SzR4 WT and mutants |  |  |  |
| --- | --- | --- | --- |
| Mutation type | Mutation | $\lambda_{\max}$ (nm) | $\Delta\lambda_{\max}$ (nm) |
| Mutation to SzR2-type amino acid | WT | 555 | 0 |
|  | N100M | 544 | -11 |
|  | V103I | 551 | -4 |
|  | F122L | 547 | -8 |
|  | S125Y | n. d. | n. d. |
|  | W161F | 553 | -2 |
| Mutation to SzR3-type amino acid | N100T | 551 | -4 |
|  | V103T | n. d. | n. d. |
| Mutation of P158 and T187 | P158T | 553 | -2 |
|  | T187V | n. d. | n. d. |

n. d.: not determined
| SzR2 WT and mutants |  |  |  |
| --- | --- | --- | --- |
| Mutation type | Mutation | $\lambda_{\max}$ (nm) | $\Delta\lambda_{\max}$ (nm) |
| Mutation to SzR4-type amino acid | WT | 542 | 0 |
|  | M101N | 543 | 1 |
|  | I104V | 545 | 3 |
|  | L121F | 541 | -1 |
|  | Y124S | 536 | -6 |
|  | F164W | 543 | 1 |

| SzR3 WT and mutants |  |  |  |
| --- | --- | --- | --- |
| Mutation type | Mutation | $\lambda_{\max}$ (nm) | $\Delta\lambda_{\max}$ (nm) |
| Mutation to SzR4-type amino acid | WT | 540 | 0 |
|  | T103N | 543 | 3 |
|  | T106V | 538 | -2 |

To determine the residues responsible for the color tuning, we comprehensively swapped the residues around the β-ionone ring between SzR4-SzR2 and SzR4-SzR3 and measured the *λ*_max_ values of the swapped mutants (Table 2 and Extended Data Fig. 4). The mutants of SzR4 to SzR2-type (SzR4 N100M) and SzR3-type (SzR4 N100T) showed 11- and 3-nm blue-shifts, respectively. By contrast, 1- and 3-nm red-shifted absorptions were observed for SzR2 M101N and SzR3 T103N. These results suggest that the difference in the amino acid at the SzR4 N100 position is one of the color tuning factors, as in type-1 rhodopsins. Moreover, the mutation of SzR4 V103 to the SzR2-type residue (I) induced a 4-nm blue shift, while the *λ*_max_ of SzR2 I104V was 3-nm longer as compared to SzR2 WT. Hence, V103 near the β-ionone ring in SzR2 also contributes to the absorption difference from SzR4. A methionine is present at this position in BR (M118) and most type-1 rhodopsins. The mutation of this residue to a smaller amino acid allows the rotation of the C6-C7 bond of retinal, connecting the β-ionone and polyene chain, and causes blue-shifted *λ*_max_ values in channelrhodopsin (C1C2) and archaerhodopsin-3^20^. This result suggests that the replacement of the smaller valine with the larger isoleucine at this position in SzRs would generate blue-shifted *λ*_max_ values, as in type-1 rhodopsins.

Overall, the structure-based mutagenesis study demonstrated that the amino-acid differences in N100 and V103 are essential factors for color tuning in SzRs. N100 and V103 are conserved in 61% and 82% of the 85 SzR homologs (Fig. 4b), and are less conserved as compared to the other residues in the retinal binding site. Thus, these differences create the diversity of the absorption spectra in SzRs.

SzR4 P158 in TM5 and T187 in TM7 are highly conserved in 96% and 100% of the SzRs (Fig. 4b), and the homologous residues in type-1 rhodopsins play a color tuning role^21,22^. Mutating the former to threonine or the latter to alanine makes *λ*_max_ longer for many type-1 rhodopsins. To determine whether these color tuning rules also apply in SzR4, we constructed the SzR4 P158T and T184A mutants. SzR4 P158T showed a 2-nm shorter *λ*_max_ than that of SzR4 WT (Table 2 and Extended Data Fig. 4), suggesting that the color tuning rule at this position is different between SzR and type-1 rhodopsins. By contrast, SzR4 T184A displayed a 7-nm red-shifted *λ*_max_ as compared to that of SzR4 WT. A similar red-shift by the mutation of an –OH bearing residue at the same position was reported in several type-1 rhodopsins^21,22^, and SzR4 T184 has a similar effect on the excitation energy of the retinal π-electron.

### Insight into proton transport

To investigate the mechanism of inward proton transport, we compared the SzR4 and BR structures. In the outward H^+^ pumping BR, an H^+^ is transferred from RSB to the proton acceptor D85 in the early stage of the photocycle at ~10^-5^ sec, and this H^+^ is finally released to the extracellular milieu via a proton release group, consisting of E194, E204 and a hydrated water between them^23^ (Fig. 5a and Extended Data Fig. 3c). However, in SzR4, D85 is replaced with the hydrophobic residue F70. The F70 side chain is directed toward the membrane environment and not involved in the interaction with the RSB (Fig. 5b and Extended Data Fig. 3d). Moreover, the extracellular H^+^ acceptors E194 and E204 in BR are replaced with A168 and T176 in SzR4, respectively. There is no other specific extracellular H^+^ acceptor in SzR4. The SzR4 counterion D184 forms salt bridges with the RSB and R67, maintaining the low pKa of D184 and preventing its protonation. The hydrogen bond between Y71 and D184 may block the RSB in SzR4 from interacting with D184. These structural observations prove that SzR cannot work as an outward proton pump.

**Fig. 5.**
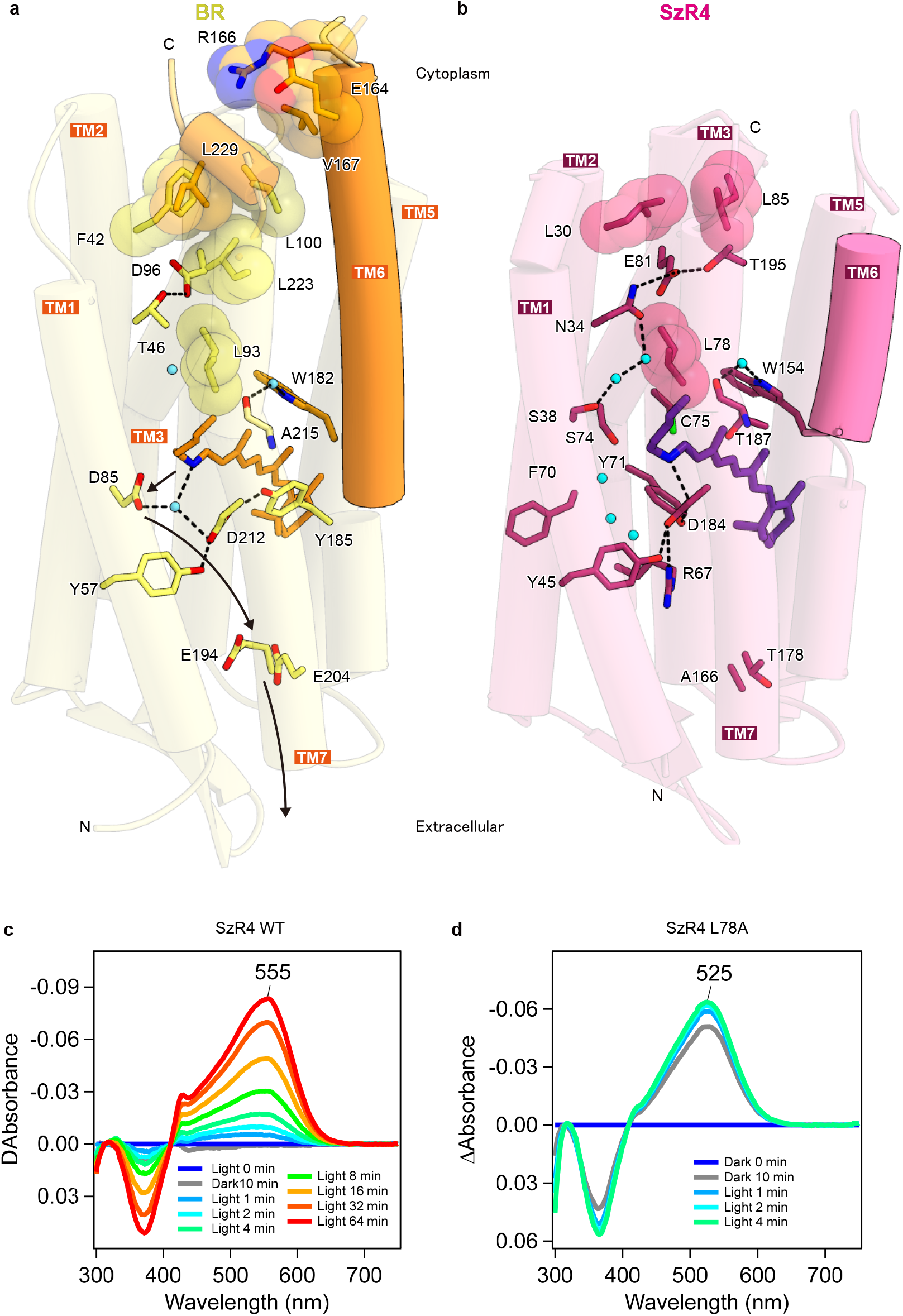
Essential residues for inward proton uptake. **a, b**, Essential residues for proton transfer in SzR4 (**a**) and BR (**b**). Waters are shown as cyan spheres. **c**, **d**, The difference absorption spectra between after and before hydroxylamine bleaching reactions of SzR4 WT (**c**) and SzR4 L78A (**d**) in solubilized *E. coli* membranes. The *λ*_max_ of each SzR and mutant was determined by the positions of the absorption peaks of the original proteins indicated in each panel, and the absorption of retinal oxime produced by the hydrolysis reaction of RSB and hydroxylamine was observed as peaks around 360–370 nm. The reaction was first performed in the dark for 10 min and then exposed to light for 64 min. Whereas no detectable bleaching of the visible region was observed for SzR4 WT in the first 10 min of the reaction in the dark, ca. 70% protein was bleached for SzR4 L78A during the same time period.

In BR, a water molecule (water402) bridges the RSB and the counterions D85 and D212 via hydrogen bonding interactions (Fig. 5a and Extended Data Fig. 3c). This strongly hydrogen-bonded water molecule is observed in all outward H^+^ pumping rhodopsins^24^ and the xenorhodopsins *Po*XeR^25^ and *Ns*XeR^16^. Around the RSB-counterion complex in SzR4, three water molecules are present in the space opened by the flipping of the F70 side chain (Fig. 5b and Extended Data Fig. 3d). These water molecules form an extensive water-mediated hydrogen-bonding network with S44, R67, S74, and the counterion D184. However, the RSB in SzR4 does not form any hydrogen-bonding interactions with the water molecules. This is in excellent agreement with the previous Fourier transform infrared (FTIR) analysis of SzR1, which indicated the presence of several water molecules around the chromophore that are not strongly hydrogen-bonded^13^. Thus, the absence of the strongly hydrogen-bonded water is a unique structural feature of SzR4, and might be associated with its function.

On the cytoplasmic side, E81 forms hydrogen bonds with N34 and T195, stabilizing its low pKa and negative charge (Fig. 5b). In BR, the equivalent residue D96 works as a cytoplasmic proton donor, supplying an H^+^ to the deprotonated RSB in the M intermediate during the outward H^+^ pump cycle^1^ (Fig. 5a). In SzRs, E81 plays a critical role in the H^+^ release process upon M-formation during the inward H^+^ pump cycle. The E81Q mutant of SzR4 lost the H^+^ transport activity, whereas the E81D mutant retained it (Extended Data Fig. 5a, b), suggesting that the negative charge of E81 plays an essential role in the H^+^ transport activity, as in SzR1.

However, a previous FTIR analysis indicated that the H^+^ is not metastably trapped by E81 in the L/M intermediate of SzR1, unlike *Po*XeR^10^. Instead, it is directly released into the cytoplasmic milieu in SzR and does not interact with the protein in the L/M state^13^. In SzR4, E81 is closer to the cytosol, since the cytoplasmic parts of TMs 2, 6, and 7 are shorter than those in the other type-1 rhodopsins, as described above (Fig. 5b and Extended Data Fig. 3a, b). E81 is separated from the solvent by only two leucine residues, L30 and L85, and easily exposed to the solvent by the light-induced structural change. An H^+^ would be attracted to the negative charge of E81 and released to the cytoplasm through the solvent water molecules.

What light-induced structural changes enable the inward H^+^ release? Recent time-resolved study of BR with millisecond time resolution (TR-SMX) has shown that the rotation of L93 opens the hydrophobic barrier between the RSB and D96 (Fig. 5a), creating space for the three water molecules that connect them^26^. This structural change allows the H^+^ transfer to the RSB. In SzR4, the equivalent residue L78 also forms the hydrophobic barrier between the RSB and E81 (Fig. 5b). Three hydrating waters exist around L78, as in BR. Thus, a similar rotation of L78 would create a water-mediated transport network from the RSB to the cytosol, with the H^+^ released to the cytoplasm through the network.

To investigate the importance of L78 for inward proton transport, we constructed the L78A mutant. No pH change was observed upon light illumination of *E. coli* cells expressing SzR4 L78A (Extended Data Fig. 5a, b), suggesting that L78 plays a critical role in the inward H^+^ transport function. Furthermore, the RSB in SzR4 WT was not hydrolyzed by hydroxylamine (HA) in the dark (Fig. 5c), whereas that in SzR4 L78A was breached by HA even without light exposure (Fig. 5d). In this mutant, the RSB would be more accessible to external solvents on the cytoplasmic side and small hydrophilic molecules such as HA. This result supports the solvent access to E81 during the photocycle and the untrapped inward H^+^ release by SzR4.

### Working model of inward proton release

We mutated the residues on a putative proton transport pathway in SzR4, and all of the mutations reduced the H^+^ transport activity (Extended Data Fig. 5a, b). The mutants of the residues in TMs 2, 4, 6, and 7 retained the transport activity itself, except for the counterion mutant (D184N), whereas those in TM3 completely lost it (S74A, C75A, C75S, C75T, L78A, and E81A). These results suggest the functional importance of TM3, in that the light-induced structural change of TM3 enables the inward proton transport.

Integrating these findings, we propose a structure-based working model of inward H^+^ release (Fig. 6). In M-rise, the protein moieties, including TM3, undergo structural changes, disrupting the hydrogen-bonding network around E81 and the two hydrophobic barriers above and below E81. Thus, a water-mediated transport network is formed between the RSB to the cytosol. Then, the RSB is deprotonated, and the H^+^ is released to the solvent trough the network, attracted by the negative charge of E81. We refer to this mechanism as untrapped inward H^+^ release.

**Fig. 6.**
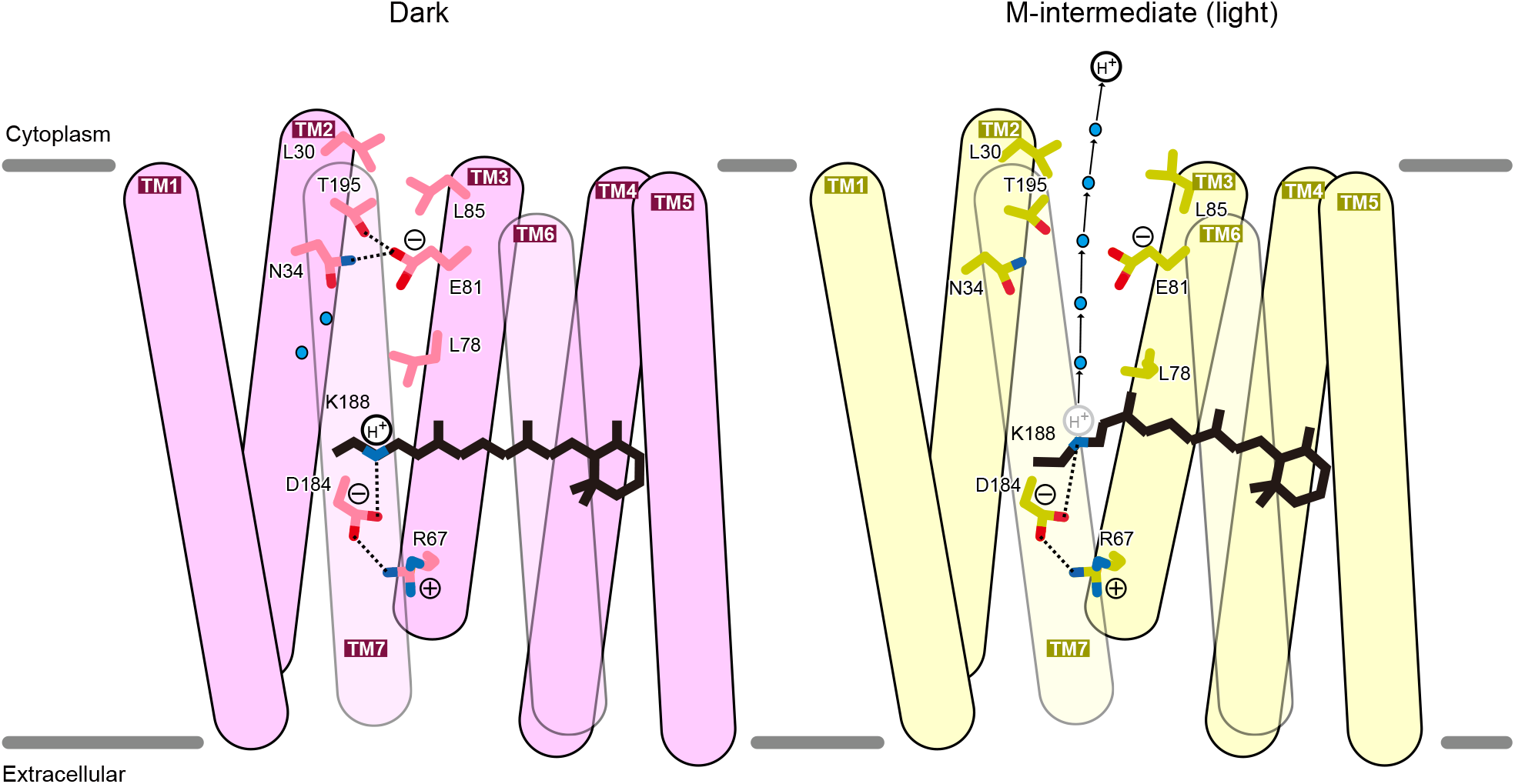
Working model of inward proton release by SzR. Models of dark and M-intermediate of SzR4. Water molecules are shown as cyan spheres, Hydrogen-bonding interactions are indicated by black dashed lines.

To inwardly uptake an H^+^, the deprotonated RSB should be re-protonated from the extracellular milieu. In *Po*XeR, the branched thermal isomerization of retinal from the *13-cis-15-anti* to *all-trans-15-anti* and *13-cis-15-syn* configurations is the rate limiting process for the reprotonation of RSB during the M-decay^10^. The *13-cis-15-anti* to all-*trans*-15-*anti* isomerization changes the inward-directed orientation of the lone pair on the nitrogen atom of RSB toward the outward-directed one, and the H^+^ can access the RSB from the extracellular side. An extensive hydrogen-bonding network, including seven hydrating water molecules, exists between the RSB and the extracellular side (Fig. 6a). Since there is no specific extracellular H^+^ donor, as suggested by the comprehensive mutations of SzR1^13^, the H^+^ is directly taken up from the extracellular milieu simultaneously with the thermal isomerization of the retinal chromophore in the M decay, through this hydrogen-bonding network as in *Po*XeR^27^.

### Comparison of SzR4 and *Ns*XeR

SzRs and XeR similarly function as inward proton pumps^13,16^, despite their distant sequence similarity. To explore their common structural features as inward proton pumps, we compared the SzR4 and *Ns*XeR structures. As described above, SzR4 has the shortest TM6, and it enables the “untrapped” inward proton release. However, TMs 2, 5, and 6 of *Ns*XeR are longer than those of SzR4 by over two turns (Fig. 6a, b and Extended Data Fig. 6a, b). Unlike SzR4, the N-terminus and ICL1 in *Ns*XeR contain characteristic α-helices. SzR4 superimposes well on BR (R.M.S.D. =1.22 Å), rather than *Ns*XeR (R.M.S.D. =1.51 Å). At the secondary structure level, SzR4 and *Ns*XeR do not share common structural features as inward proton pumps.

We also compared the proton transport pathways in SzR4 and *Ns*XeR (Fig. 6a, b). F70 and D184 in SzR4 are replaced with D76 and P209 in *Ns*XeR, respectively. Thus, SzR4 and *Ns*XeR similarly have a single counterion, although its relative position in the structure is different. The smaller negative charge on the extracellular side of the RSB, as compared to the outward H^+^ pumps with double counterions, would increase the directionality of H^+^ transfer to the cytoplasmic H^+^ acceptor relative to the extracellular counterion. *Ns*XeR has the proton acceptors H48 and D220 on the cytoplasmic side, and the H^+^ is trapped by these residues in the M state. However, these residues are not conserved in SzR4, and the H^+^ is not trapped in the M state. Moreover, the other residues on the proton transport pathway are not conserved between SzR4 and *Ns*XeR, suggesting that they have entirely different inward H^+^ release mechanisms. Nevertheless, SzR4 and *Ns*XeR similarly have the extensive water-mediated hydrogen-bonding network in their extracellular halves, and the H^+^ is easy to access from the extracellular milieu to the RSB.

## Discussion

Our SzR4 structure offers numerous insights into the structure-function relationships and color tuning mechanisms of the SzR family members. Although SzRs are phylogenetically located at an intermediate position between type-1 rhodopsins and HeRs, SzRs are structurally similar to type-1 rhodopsins, and thus we classified SzRs with them. Since the cytoplasmic part of TM6 is the shortest among the microbial rhodopsins, E81 is located near the cytosol and plays a critical role in the inward proton transport. Given that the H^+^ is not trapped in E81, light-induced structural changes would displace the rotamer of L78, releasing the H^+^ to the solvent water molecules, attracted by the negative charge of E81 (Fig. 6).

By contrast, in AntR, the proton transport rate of E81Q is reportedly similar to that of the WT^15^, indicating that the negative charge of E81 is not essential for its function. To understand the proton uptake mechanism of AntR, we constructed a homology model of AntR based on the SzR4 structure (Extended Data Fig. 7a, b). In this model, similar to D96 in BR, E81 does not form any polar interactions and is surrounded by hydrophobic residues. Thus, E81 would be protonated and not associated with the proton transport in AntR. Notably, D30 forms a hydrogen-bonding network with R84 and Y193, which are located somewhat closer to the cytoplasmic side as compared with E81. These residues are unique in AntR (Extended Data Fig. 7a). A previous study demonstrated that the R84A mutant retains the proton transport activity, whereas M-formation is absent in the photocycle of the Y193F mutant. These observations suggest that the hydrogen-bonding interaction between D30 and Y193 is critical in the photoactivation of AntR. Instead of E81, D30 might be deprotonated and negatively charged, and thus play an essential role in the inward proton release by AntR. The homology model of AntR represents the diversity of the inward proton transport mechanisms among the SzR/AntR family members.

**Fig. 7.**
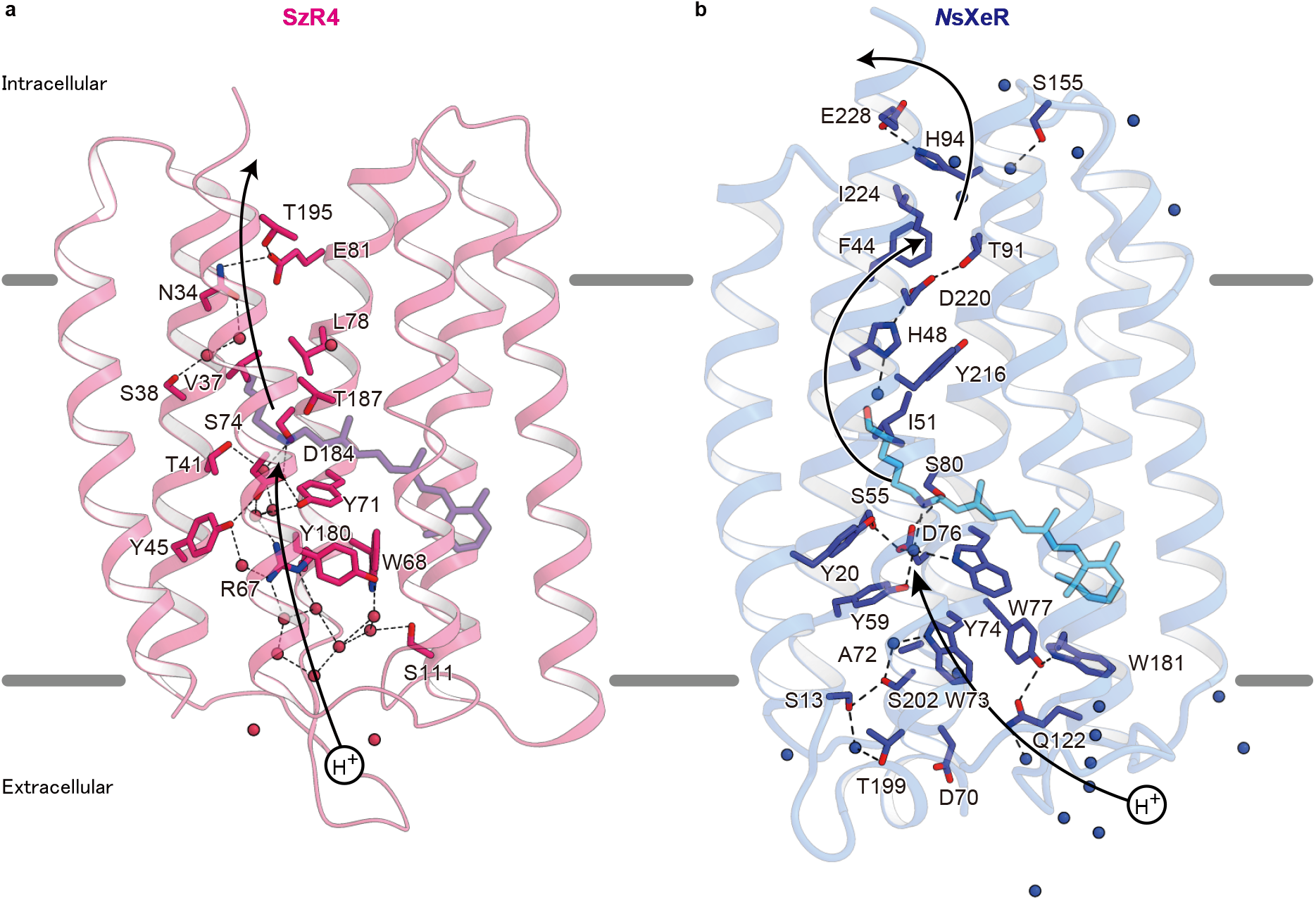
Ion translocation pathway in inward H^+^ pumps. **a, b**, Putative key residues inside SzR4 (**a**) and *Ns*XeR (**b**) are shown. Black arrows show the proposed proton path.

## Acknowledgements

The diffraction experiments were performed at SPring-8 BL32XU (proposals 2018B2544 and 2019A0153). We thank the members of the Nureki lab and the beamline staff at BL32XU of SPring-8 (Sayo, Japan) for technical assistance during data collection; Drs. O. Béjà, R. Ghai, S. P. Tsunoda and S. Hososhima for useful discussions; and R. Nakamura and Y. Yamauchi for technical assistance. This research was partially supported by the Platform Project for Supporting Drug Discovery and Life Science Research (Basis for Supporting Innovative Drug Discovery and Life Science Research (BINDS)) from AMED under grant number JP19am0101070 (support number 1627). This work was supported by JSPS KAKENHI grants 16H06294 (O.N.), 20H05437, 20K15728 (W.S.), 25104009, 15H02391, 18H03986 (H.K.), and 17H03007 (K.I.), and by JST PRESTO (JPMJPR15P2) and CREST (JPMJCR1753 and JPMJCR17N5).

## Author contributions

A.H. screened the homologs of the SzRs and crystallized SzR4. A.H. and W.S. solved and refined the structure. M.K. and K.I. performed the functional analyses of SzRs, with the supervision by H.K. T.I. performed the homology modeling of AntR. The manuscript was mainly prepared by A.H., W.S., K.I., and O.N. The research was supervised by W.S., K.I., and O.N. The authors declare no competing financial interests. Coordinates and structure factors have been deposited in the Protein Data Bank, under the accession number XXXX. The X-ray diffraction images are also available at SBGrid Data Bank (https://data.sbgrid.org/), under the ID YYY.

## Methods

### Expression and purification

The gene encoding SzR4 (GenBank ID: TFG21677.1), with codons optimized for an *E. coli* expression system, was synthesized (Genscript) and subcloned into the pET21a(+)-vector with an N-terminal 6×His-tag. The protein was expressed in *E. coli* C41(Rosetta). Protein expression was induced by 1 mM isopropyl β-D-thiogalactopyranoside (IPTG) for 20 h at 25 °C, and then the culture was supplemented with 10 μM *all-trans* retinal (Sigma Aldrich). The harvested cells were disrupted by sonication in buffer, containing 20 mM Tris-HCl (pH 7.5), 20% glycerol. The crude membrane fraction was collected by ultracentrifugation at 180,000g for 1 h. The membrane fraction was solubilized for 1 h at 4 °C, in buffer, containing 20 mM Tris-HCl (pH 7.5), 150 mM NaCl, 1% DDM, and 10% glycerol. The supernatant was separated from the insoluble material by ultracentrifugation at 180,000g for 20 min, and incubated with TALON resin (Clontech) for 30 min. The resin was washed with ten column volumes of buffer, containing 20 mM Tris-HCl (pH 7.5), 500 mM NaCl, 0.03% DDM, and 15 mM imidazole. The protein was eluted in buffer, containing 20 mM Tris-HCl (pH 7.5), 500 mM NaCl, 0.03% DDM, and 200 mM imidazole. The eluate was concentrated and loaded onto a Superdex200 10/300 Increase size-exclusion column, equilibrated in buffer, containing 20 mM Tris-HCl (pH 7.5), 150 mM NaCl and 0.03% DDM. Peak fractions were pooled, concentrated to 30 mg ml^-1^ using a centrifugal filter device (Millipore 50 kDa MW cutoff), and frozen until crystallization.

### Crystallization

The protein was reconstituted into monoolein at a weight ratio of 1:1.5 (protein: lipid). The protein-laden mesophase was dispensed into 96-well glass plates in 30 nL drops and overlaid with 800 nL precipitant solution, using a Gryphon robot (ARI). Crystals of SzR4 were grown at 20°C in precipitant conditions containing 20% PEG500DME, 100 mM Na-acetate, pH 4.75, and 250 mM MgSO4. The crystals were harvested directly from the LCP using micromounts (MiTeGen) or LithoLoops (Protein Wave) and frozen in liquid nitrogen, without adding any extra cryoprotectant.

### Data collection and structure determination

X-ray diffraction data were collected at the SPring-8 beamline BL32XU with an EIGER X 9M detector (Dectris), using a wavelength of 1.0 Å. In total, 148 small-wedge (10° per crystal) datasets were obtained using a 15×10 μm^2^ beam. The collected images were processed with KAMO^28^. Each data set was indexed and integrated with XDS^29^ and then subjected to a hierarchical clustering analysis based on the unit cell parameters, using BLEND. After outlier rejection, 127 datasets were finally merged with XSCALE^29^. The SzR4 structure was determined by molecular replacement with PHASER^30^, using the structure of bacteriorhodopsin (PDB code: 1M0K)^31^. Subsequently, the model was rebuilt and refined using COOT^32^ and phenix.refine^33^. Figures were prepared with CueMol (http://www.cuemol.org/ja/).

### Laser flash photolysis

For the laser flash photolysis measurement, SzR4 was purified and reconstituted into a mixture of POPE (Avanti Polar Lipids, AL) and POPG (sodium salt, Avanti Polar Lipids, AL) (molar ratio = 3:1), with a protein-to-lipid molar ratio of 1:50, in buffer, containing 20 mM Tris-HCl (pH 8.0) and 100 mM NaCl. The absorption of the protein solution was adjusted to 0.8–0.9 (total protein concentration ~0.25 mg ml^-1^) at an excitation wavelength of 532 nm. The sample was illuminated with a beam of the second harmonics of a nanosecond-pulsed Nd^3+^-YAG laser (*λ* = 532 nm, 3 mJ pulse^-1^, 1 Hz) (INDI40; Spectra-Physics, CA). The transient absorption spectra were obtained by monitoring the intensity change of white-light from a Xe-arc lamp (L9289-01, Hamamatsu Photonics, Japan) passed through the sample, with an ICCD linear array detector (C8808-01, Hamamatsu, Japan). To increase the signal-to-noise (S/N) ratio, 90 spectra were averaged and a singular-value-decomposition (SVD) analysis was applied. To measure the time-evolutions of transient absorption changes at specific wavelengths, the light from the Xe-arc lamp (L9289-01, Hamamatsu Photonics, Japan) was monochromated with a monochromater (S-10, SOMA OPTICS, Japan), and the change in the intensity after photo-excitation was monitored with a photomultiplier tube (R10699, Hamamatsu Photonics, Japan) equipped with a notch filter (532 nm, bandwidth = 17 nm, Semrock, NY) to remove the scattered pump pulses. To increase the S/N ratio, 100 signals were averaged.

### Measurement of absorption maximum wavelength by hydroxylamine bleaching

The *λ*_max_ values of the wildtype and mutants of SzR4, SzR2, and SzR3 were determined by bleaching the protein with hydroxylamine, according to the previously reported method^22^. *E. coli* cells expressing rhodopsins were washed three times with buffer, containing 50 mM Na2HPO4 (pH 7) and containing 100 mM NaCl. The washed cells were treated with 1 mM lysozyme for 1 hr at room temperature, and then disrupted by sonication. To solubilize the rhodopsins, 3% DDM was added and the samples were stirred overnight at 4 °C. The rhodopsins were bleached with 500 mM hydroxylamine in the dark or under yellow light illumination (*λ* > 500 nm) from the output of a 1 kW tungsten-halogen projector lamp (Master HILUX-HR, Rikagaku) passed through a glass filter (Y-52, AGC Techno Glass). The absorption change upon bleaching was measured by a UV-visible spectrometer (V-730, JASCO, Japan) equipped with an integrating sphere (ISV-922, JASCO, Japan).

### Homology modeling of AntR

The AntR homology model was built based on the crystal structure of SzR using Modeller^34–37^. The input sequence alignment was generated from the sequence of SzR (residues 1-199) and AntR.

## Extended Data Figures

**Extended Data Fig. 1. Structural analysis of SzR4.**

**a, b**, Structural comparisons of molA with molB (**a**) and molC (**b**). **c-e**, Oligomeric interface of the SzR4 structure. **f**, Conservation of the trimer interface in the SzR4 structure. The sequence conservation among 85 SzRs was calculated using the ConSurf server (http://consurf.tau.ac.il) and colored from cyan (low) to maroon (high).

**Extended Data Fig. 2. Structure-based alignment of SzRs.**

Multiple amino acid sequential alignments of SzR4 with typical microbial rhodopsins. The amino acid sequences were aligned using ClustalW^38^. SzR1, SzR2, SzR3, SzR TE_8_00242, SzR_TE_5S_00009, SzR un_Tekir_02407, Bacteriorhodopsin (BR), *Natronomonas pharaonis* halorhodopsin (*Np*HR), green-absorbing proteorhodopsin (GPR), *Krokinobacter* rhodopsin 2 (KR2), *Parvularcula oceani* xenorhodopsin (*Po*XeR), *Chlamydomonas reinhardtii* channelrhodopsin 2 (*Cr*ChR2), and HeRs (HeR-48C12 and *Thermoplasmatales* archaeon SG8-52-1 heliorhodopsin (*T*aHeR)) were aligned with the sequence of SzR4. The residue numbers of SzR4 and BR are shown on top of the residues, and the positions of transmembrane helices and β-strands, based on the X-ray crystallographic structures of SzR4 and BR (PDB ID: 1M0L^31^), are indicated by rectangles and arrows, respectively. For clarity, the diverse, long N- and C-termini and interhelical loops of *Np*HR, GPR, KR2, *Cr*ChR2, *Po*XeR, *T*aHeR, and HeR 48C12 were omitted

**Extended Data Fig. 3. Comparison of SzR4 with other microbial rhodopsins.**

**a, b**, Superimposition of SzR4 with the proton-pumping rhodopsins (**a**) and the other microbial rhodopsins (**b**), determined to date. SzR4 and the other rhodopsins are colored magenta and blue, respectively. **c, d**, Comparison of the water-mediated hydrogen-bonding networks around the RSBs in BR (**c**) and SR4 (**d**).

**Extended Data Fig. 4. Determination of *λ*_max_ of SzR and mutants.**

Difference absorption spectra between after and before hydroxylamine bleaching reactions of SzR and their mutants in solubilized *E. coli* membranes. The *λ*_max_ of each SzR and mutant was determined by the positions of the absorption peaks of the original proteins indicated in each panel, and the absorption of retinal oxime produced by the hydrolysis reaction of RSB and hydroxylamine was observed as peaks around 360–370 nm.

**Extended Data Fig. 5. Inward proton transport activity of SzR4 mutants.**

**a**, pH changes upon light illumination of suspensions of *E. coli* cells expressing SzR4 wildtype and mutants, without (blue lines) and with (green lines) 10 μM CCCP. Light illumination occurred in the time regions indicated by yellow lines. **b**, Proton uptake rates of SzR4 wildtype and mutants, calculated by dividing the proton concentration change per second upon light illumination by the protein concentration.

**Extended Data Fig. 6. Structural comparison of SzR4 and *Ns*XeR.**

**a**, Superimposition of the SzR4 and *N*sXeR structures. **b**, Individual TM helices are shown after superimposition of the two rhodopsins, as in (**a**).

**Extended Data Fig. 7. Comparison of SzR4 and AntR.**

**a**, Amino acid alignment of SzR4 and AntR. **b**, The homology model of AntR.

