## supplementary file for "Crystal structure of schizorhodopsin reveals mechanism of inward proton pumping"

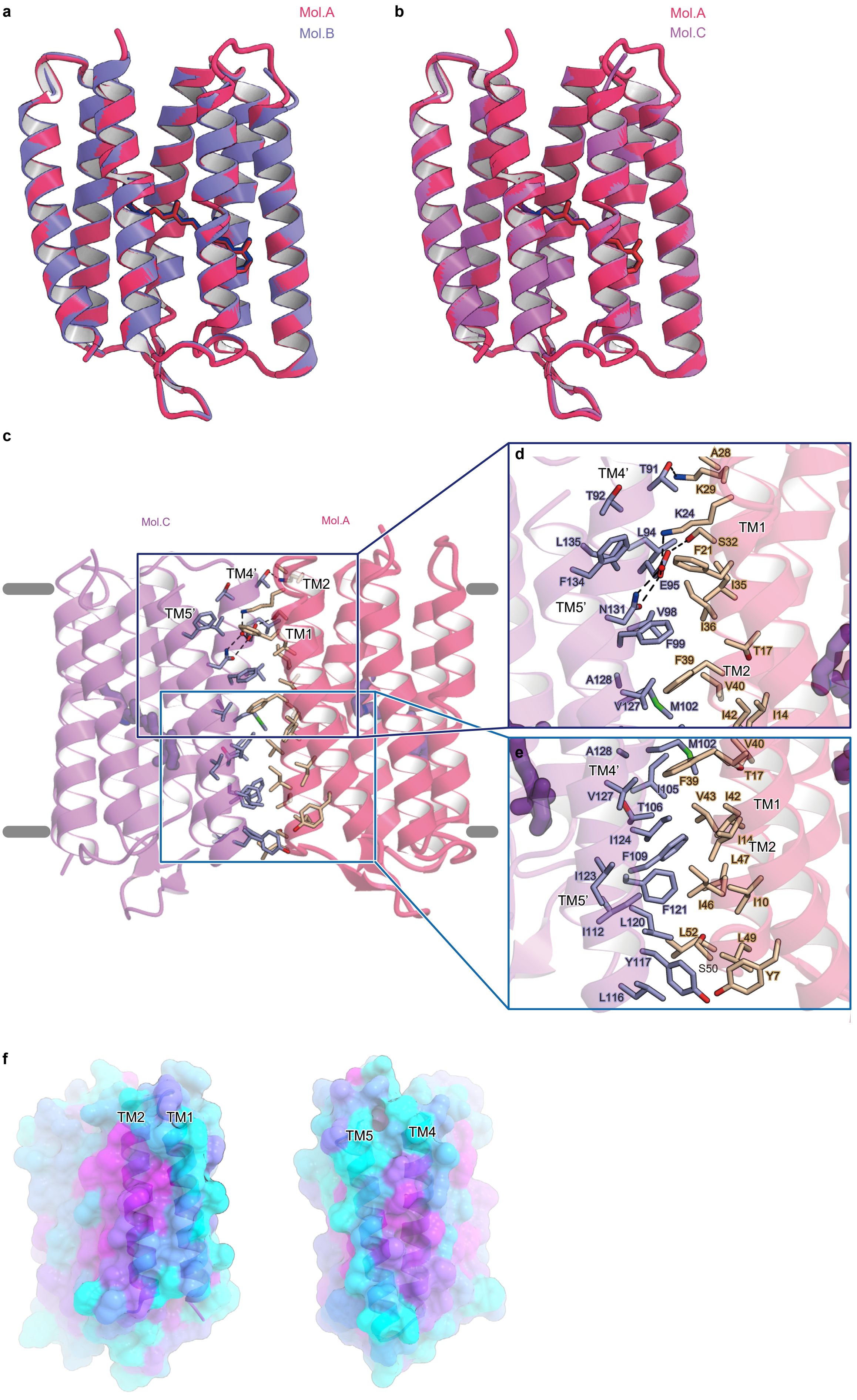


### Higuchi et at., Extended Data Figure 1

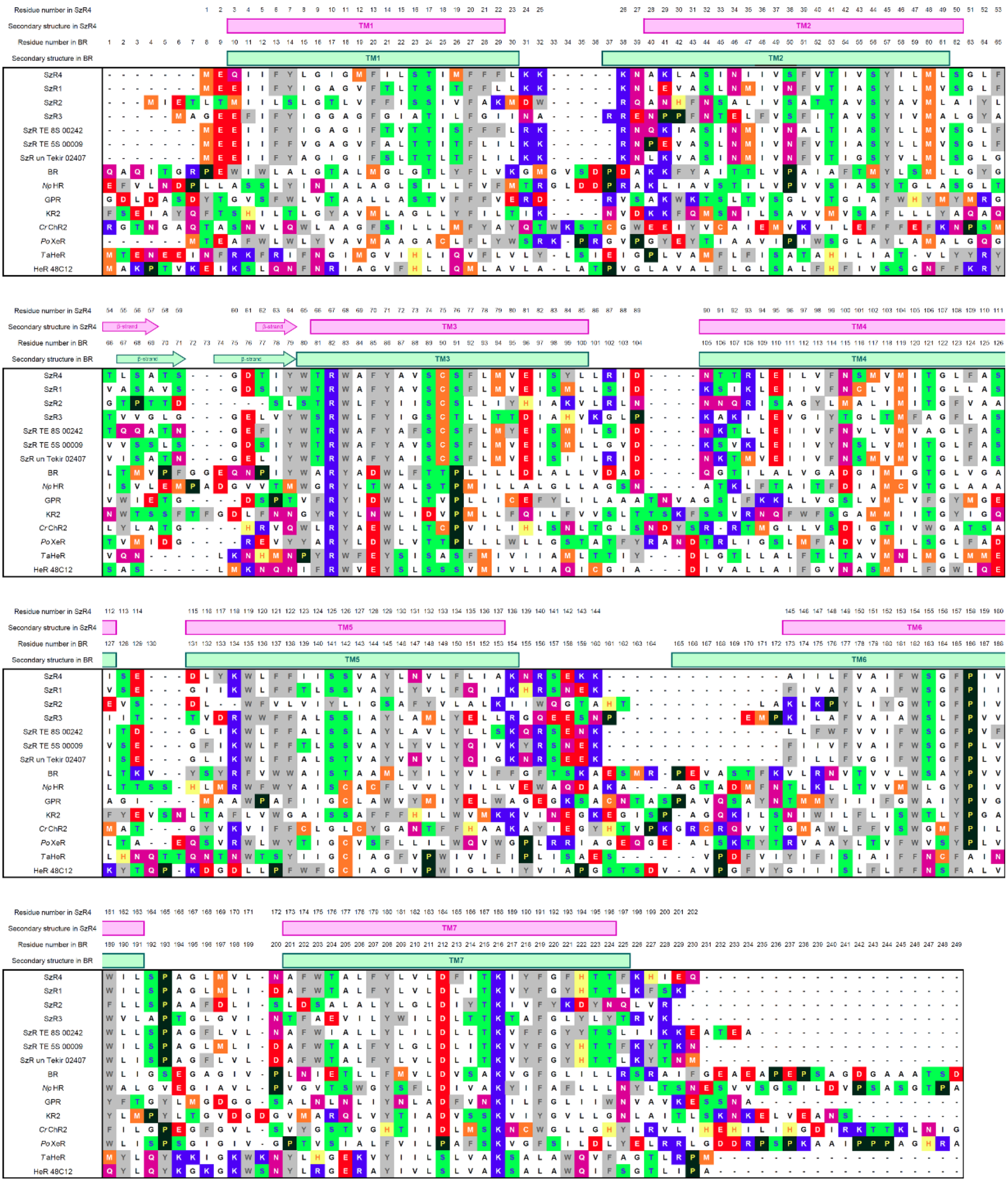


### Higuchi et at., Extended Data Figure 2

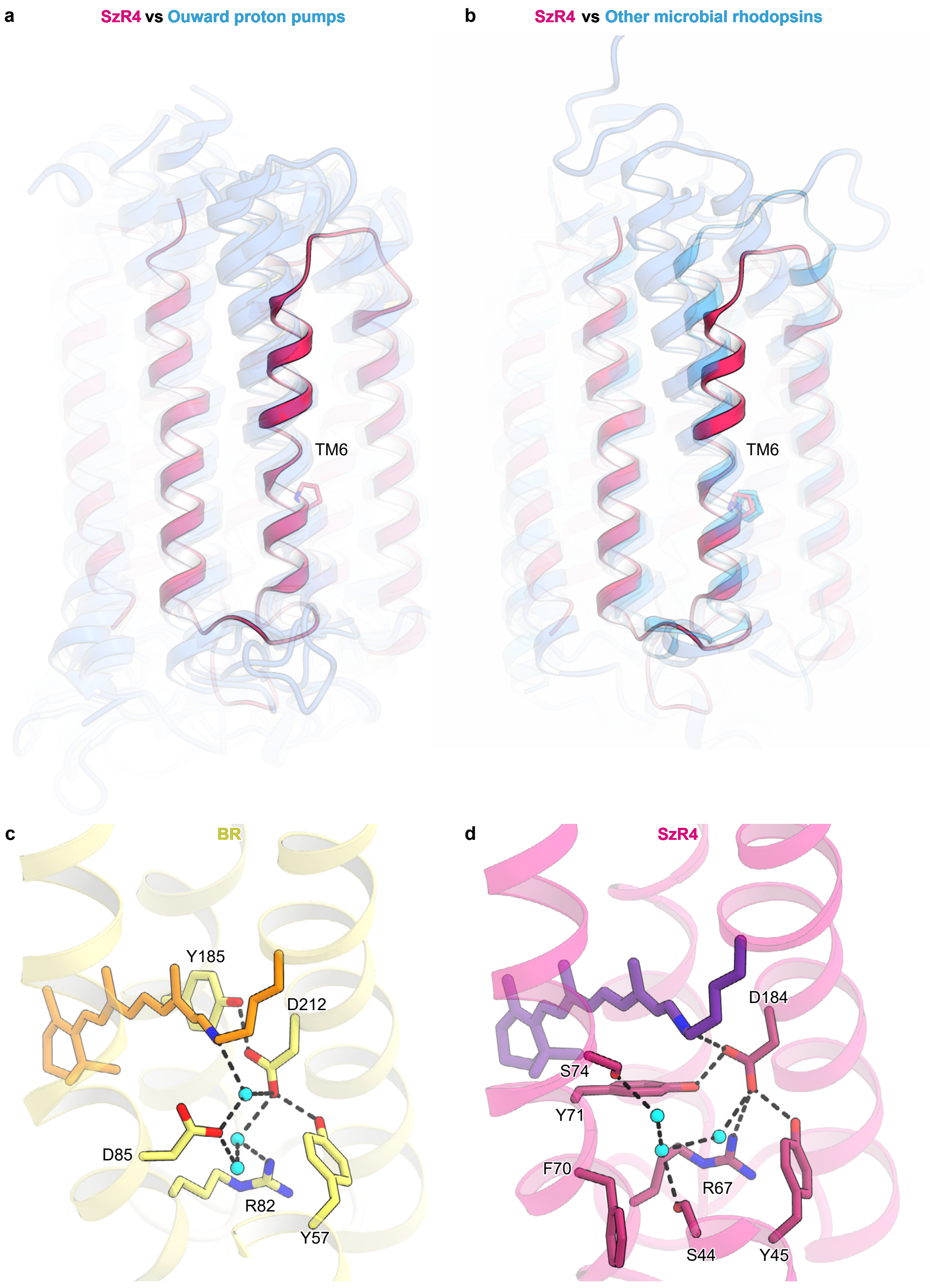


### Higuchi et at., Extended Data Figure 3

SzR4 WT and mutants

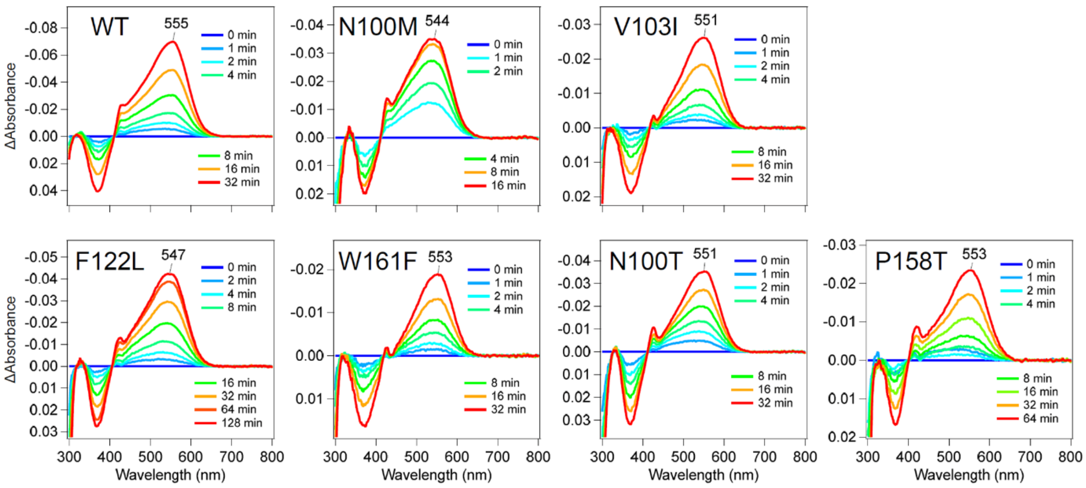


SzR2 WT and mutants

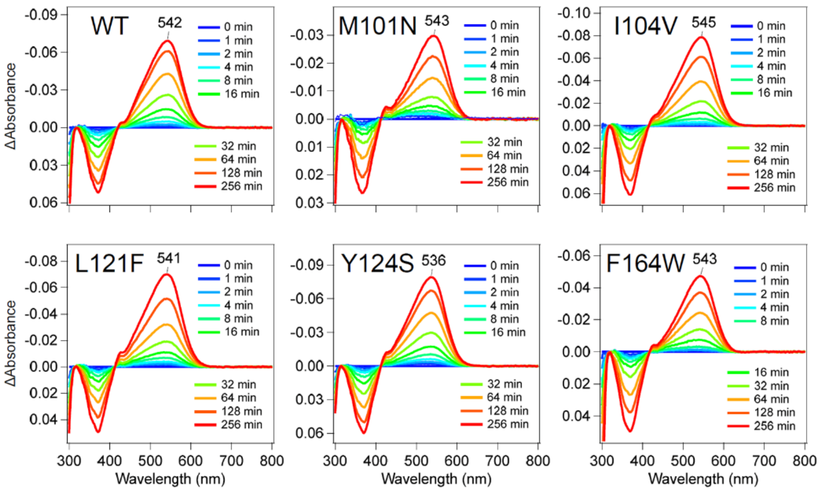


SzR3 WT and mutants

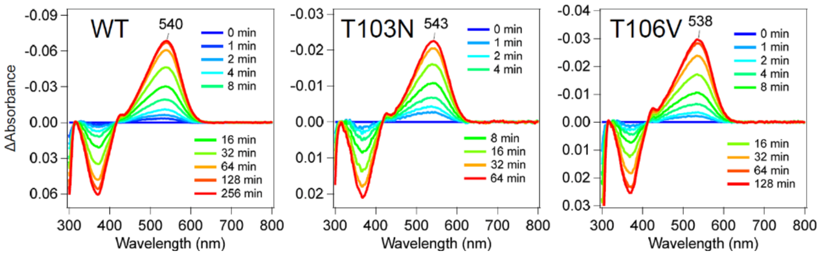


### Higuchi et at., Extended Data Figure 4

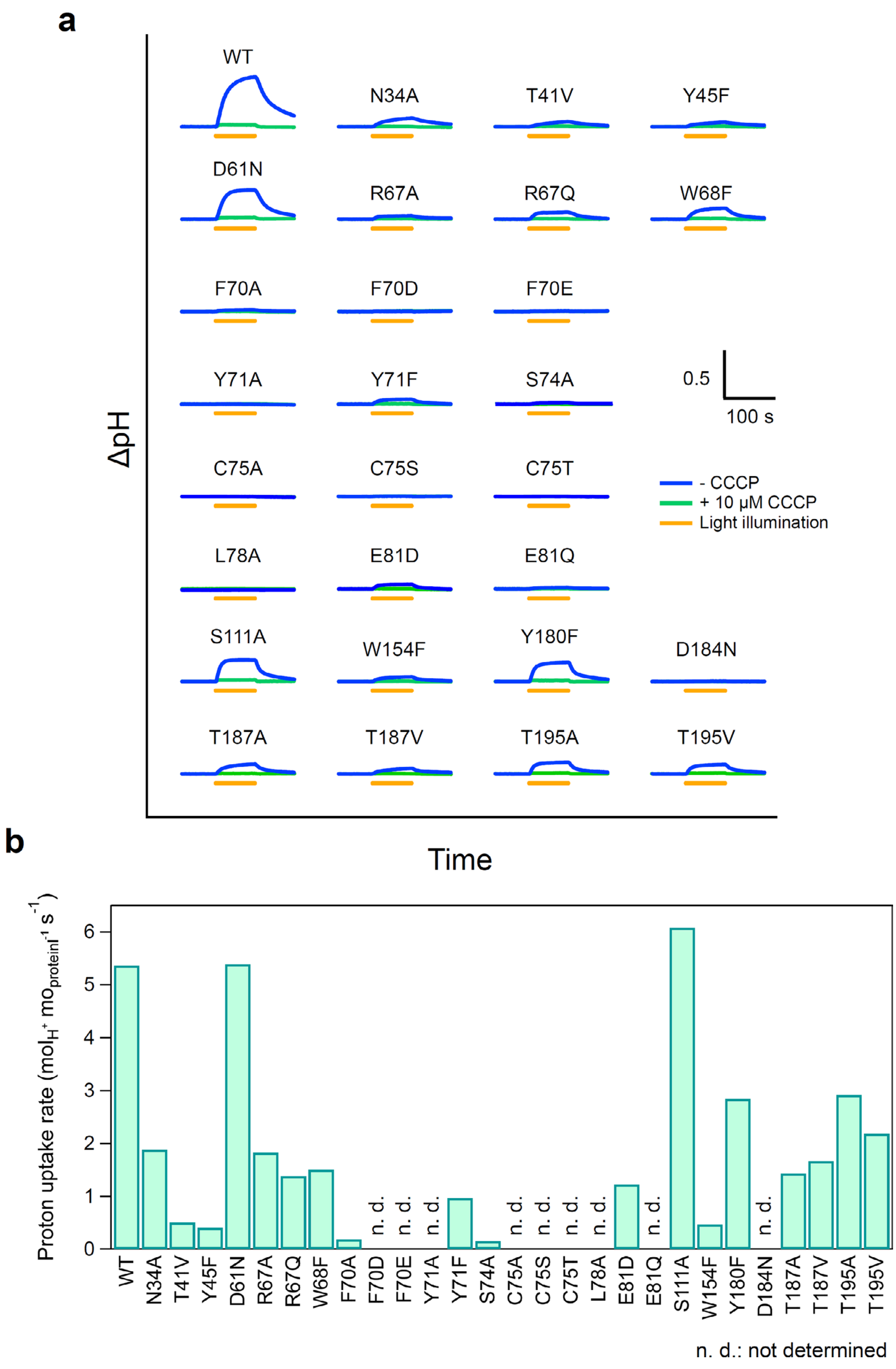


**Higuchi et at., Extended Data Figure 5**

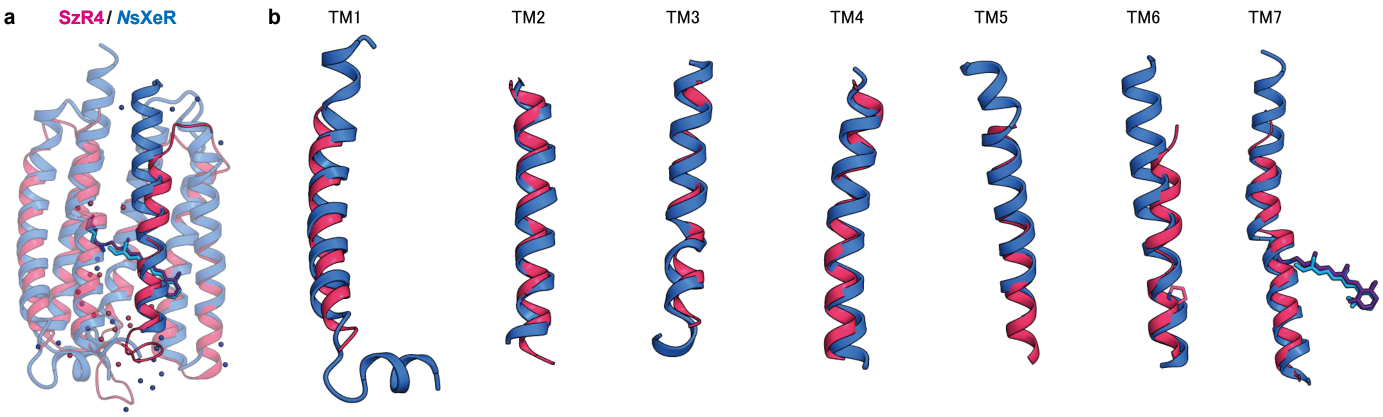


**Higuchi et at., Extended Data Figure 6**

a

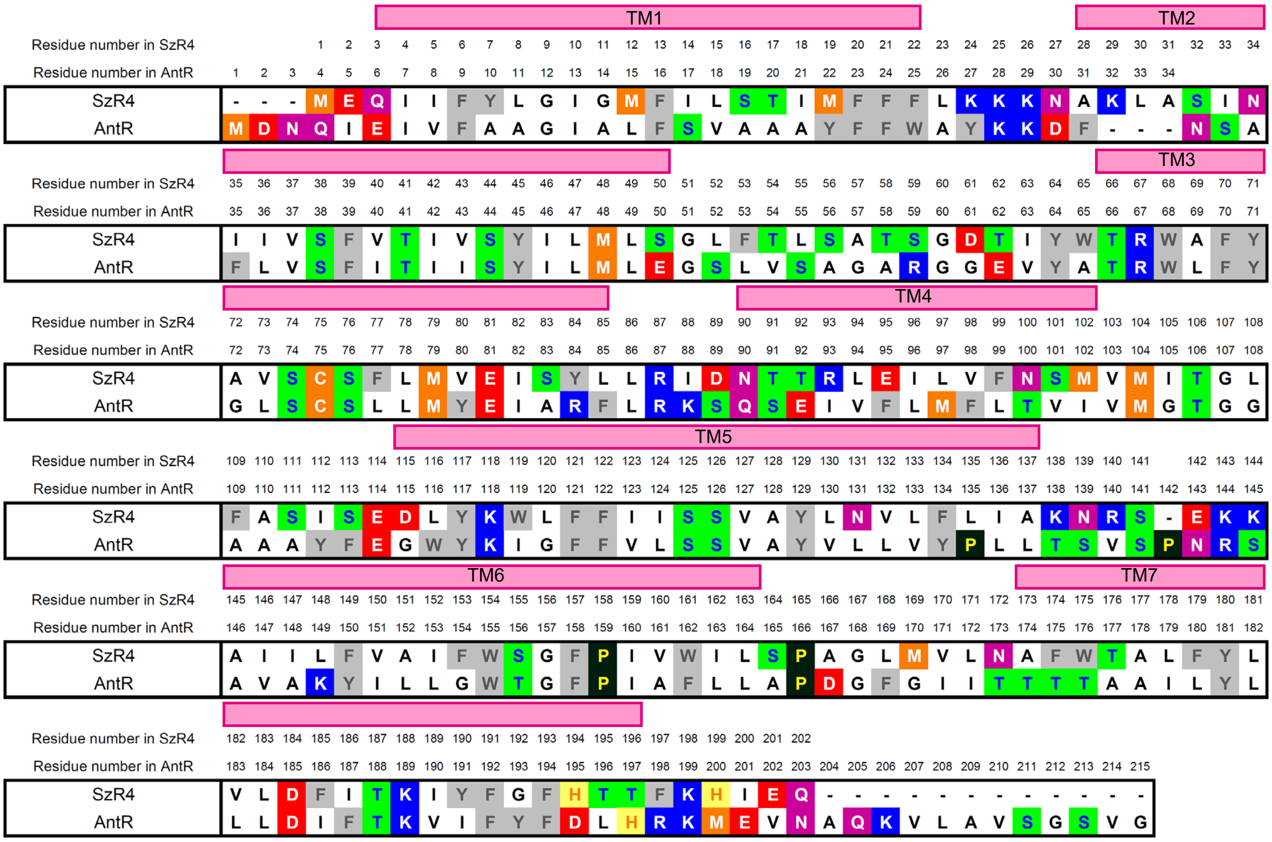


b

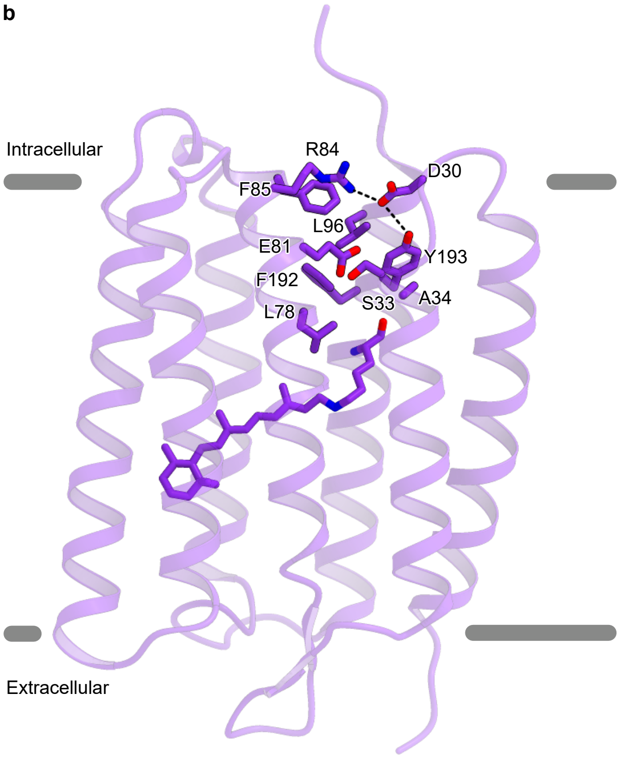


### Higuchi et at., Extended Data Figure 7
